# Survivin (BIRC5) peptide vaccine in the 4T1 murine mammary tumor model: a potential neoadjuvant T-cell immunotherapy for triple negative breast cancer

**DOI:** 10.1101/2022.07.25.501455

**Authors:** Scott R Burkholz, Charles V Herst, Richard T Carback, Paul E Harris, Reid M Rubsamen

## Abstract

A triple negative breast cancer (TNBC) model using the murine 4T1 tumor cell line was used to explore the efficacy of an adjuvanted survivin (SVN)-peptide microparticle vaccine using tumor growth as the outcome metric. We first performed tumor cell dose titration studies to determine a tumor cell dose that resulted in sufficient tumor takes but minimal morbidity/mortality within the required study period. Later, in a second cohort of mice, the SVN-peptide microparticle vaccine was administered via intraperitoneal injection at study start with a second dose given 14 days later. An orthotopic injection of 4T1 cells into the mammary tissue was performed on the same day as the administration of the second vaccine dose. The mice were followed for up to 41 days with subcutaneous measurements of tumor volume made every 3-4 days. Vaccination with SVN peptides was associated with a peptide antigen-specific γIFN Elispot response in the murine splenocyte population but was absent from the control microparticle group. At end of study, we found that vaccination with adjuvanted SVN-peptide microparticles resulted in statistically significant slower primary tumor growth rates in BALB/c mice challenged with 4T1 cells relative to the control peptide-less vaccination group. These studies suggest that T-cell immunotherapy specifically targeting survivin might be an applicable neoadjuvant immunotherapy therapy for TNBC. More pre-clinical studies and clinical trials will be needed to explore this concept further.

## 1. Introduction

The survivin (SVN), also known as BIRC5, protein is an inhibitor of apoptosis and is overexpressed in many malignancies, including breast cancer stem cells and breast tumor tissue [1], relative to adjacent normal adult cells and tissues (reviewed in [2] and [3]). These observations suggest that SVN might be an ideal tumor cell target. In fact the NCI declared SVN as a research priority more than twenty years ago[4]. Since then, SVN as a target of humoral and cellular immunity has been studied in animal models and in cancer patients. The structure of many immunogenic B cell and T cell epitopes of SVN have been characterized [2,5-8] and applied to the development of synthetic survivin-peptide cancer vaccines. These studies have provided proof of principle data supporting several currently active human immunotherapy/vaccine clinical trials targeting SVN in glioblastoma, neuroendocrine tumors, ovarian cancer, and hormone receptor positive breast cancer (e.g., NCT05163080, NCT02334865, NCT04895761). In glioblastoma, immunotherapy targeting survivin has provided significant prolongation of overall survival compared with standard radiation therapy and chemotherapy [9].

Estrogen, progesterone receptor negative, HER2 “negative” breast cancer (TNBC) accounts for up to 15%-20% of all breast malignancies, has a generally worse prognosis and fewer treatment options relative to more common forms of breast cancer. Current standard of care treatment options for recurrent TNBC have been limited to conventional cytotoxic chemotherapeutic agents due the lack of expression of molecular targets found on the more common types of breast malignancies. Adjuvant immunotherapy for TNBC is currently being explored and the approaches fall into several diverse categories (reviewed in [10]). One approach, recently approved by the FDA, combines chemotherapy with immune checkpoint inhibitors (ICI) with the goal of boosting existing adaptive immune response to breast tumor antigens. However, immunohistochemistry studies show that PD-L1 expression, one target of ICI therapy, is limited to about 20% of TNBC [11]. Clinical and histochemical data has confirmed that efficacy of anti PD-L1 mAb ICI as neoadjuvant and adjuvant treatment for TNBC patients whose tumors and/or stromal cells (including tumor infiltrating lymphocytes) express PD-L1. While the PD-L1 positive subgroup show better overall survival, the PD-L1 negative subgroup faired equally poorly, regardless of anti-PD-L1 treatment[12], suggesting that additional approaches to adjuvant immunotherapy need to exploration.

Adjuvant vaccine-based immunotherapy of TBNC has been recently reviewed [13]. Out of 42 clinical studies reviewed, only one limited exploratory human study[14] has targeted SVN in TBNC. With the goal of developing novel immunotherapies for TNBC, we turned to the 4T1 murine model of TNBC [15] to evaluate the therapeutic potential of a synthetic SVN peptide based microparticle vaccine for adjuvant immunotherapy of TNBC. We previously developed an adjuvanted, synthetic peptide - poly (lactic-co-glycolic acid) (PLGA) copolymer microparticle vaccine platform [16] capable of inducing robust therapeutic CD8+ cytotoxic T lymphocyte responses to viral antigens presented by MHC Class I molecules in non-human primate [17] and murine models [18]. In this preclinical study, we report on the effects of vaccination with adjuvanted survivin-peptide loaded microspheres on orthotopically implanted tumor growth in a 4T1 murine model of TBNC.

## 2. Materials and Methods

### 2.1. Characterization of survivin expression in 4T1 cells and normal mouse mammary tissue

To fully characterize SVN in the TNBC model mouse 4T1 cell line (ATCC, Manassas, VA), frozen 4T1 cell samples were sent to Complete Genomics (Beijing, China) for sequencing. DNA was extracted from the snap-frozen cryopreserved 4T1 cell line, and BALB/c mouse tails. Sample DNA was prepared using the Agilent SureSelect XT Mouse All Exon Kit (Agilent Technologies, Santa Clara, CA). RNA was extracted from the frozen 4T1 cell line and from also from snap-frozen, normal mammary tissue samples harvested from BALB/c mice. RNA samples were prepared for mRNA sequencing via poly-A tail capture with MGI Tech Company reagents (MGI, Shenzhen, China). Samples were sequenced by Complete Genomics on the BGISEQ-500 (BGI, Beijing, China) at 100 base pair paired end reads. DNA read coverage for normal tissue and tumor was 100x and 300x respectively, with RNA read count at 80 million paired end reads. To process the data, FASTP was used to perform quality control and adapter trimming and BWA-MEM aligned reads to the GRCm38 reference mouse genome [19-22]. The germline sequences were confirmed to match the peptides identified for vaccine administration. RNA expression of survivin was examined by adapter trimming and quality control with FASTP [19], followed by pseudoalignment with Kallisto on the GENCODE v25 mouse transcriptome for quantification in Sleuth expressed as transcripts per million [21].

### 2.2. Vaccine Design/Peptide Selection

The overall strategy and rationale for the selection of synthetic peptides used to stimulate potential CTL immune responses to SVN targets presented by murine 4T1 cells has been previously described[16,18,23]. Briefly, we identified survivin peptide antigens potentially capable of stimulating MHC Class I and Class II restricted, tumor-specific T-cell response in BALB/c mice. These peptide sequences were determined by reviewing previous publications describing SVN specific MHC Class I restricted T cells response using various peptide vaccine formulations [24,25]. In addition, NetMHCIIpan and NetMHCII [26] were utilized to identify QP19, a region predicted to bind to class II MHC to stimulate CD4 helper T-cells. The sequences of these murine class I H-2 K, D, and L binding peptides and murine class II H-2 I-E and I-A binding peptides are given in Table 1.

**Table 1.** Literature references and prediction tools supporting a BALB/c MHC match to each of the six peptides microencapsulated into the adjuvanted microsphere vaccine platform described here.

| Epitope Name | Peptide Sequence | MHC Class | Position | MHC Match Screening Method |
| --- | --- | --- | --- | --- |
| AL9 | ATFKNWPFL | I | 20-28 | [26,27] |
| AM9 | AFLTVKKQM | I | 85-93 | [25] |
| GI9 | GWEPDDNPI | I | 66-74 | [24,25] |
| TI9 | TAKTTRQSI | I | 127-135 | [25] |
| QP19 | QIWQLYLKNYRIATFKNWP | I/II | 8-26 | [26,28] |
| PADRE | AKFVAAWTLKAAA | 2 | N/A | [29,30] |

### 2.3. Vaccine Manufacture

A blend of PLGA microspheres, prepared as described in a previous publication [17], was manufactured containing individual synthetic GMP-grade peptides (Peptides International, Louisville, KY, USA), selected from the primary amino acid sequence of the murine survivin protein (Table 1). The adjuvanted microsphere vaccine formulation contained 3 µg/mg of survivin-specific class 1 and class 2 peptides and 0.5 µg/mg of the TLR-9 oligonucleotide agonist CpG (ODN-1018) (Trilink Biosciences, San Diego, CA, USA). The microspheres, approximately 6 microns in diameter, were delivered in a 200 µl volume of PBS / Tween 20 (0.01% v/v) injectate solution containing the TLR-4 agonist MPLA (Avanti Polar Lipids, Alabaster, AL, USA) at a concentration of 100 µg/ml.

### 2.4. Post vaccination evaluation of peripheral T cell responses

Enzyme-linked immunosorbent spot for interferon gamma (ELISpot) assays were performed with BALB/c splenocytes obtained from control and vaccinated surviving tumor bearing mice on day 41 after 4T1 tumor inoculation. Splenocytes were prepared as previously described [31]. The peptide antigens used in the ELISpot assays were the same as those used in the peptide vaccine and individually added to ELISpot wells at 10 µg/ml final concentration. ELISpot assay plates were prepared and processed as per the manufacturer’s instructions (3321-4HPT-10, Mabtech Inc., Cincinnati, OH). ELISpots were enumerated by machine (CTL S6 Entry M2, Shaker Heights, Cleveland, OH) to calculate the frequency of gamma interferon producing T-cells in the splenocyte populations.

### 2.5. Animal Immunizations

Previous studies with the adjuvanted microsphere platform have shown that T-cell expansion capable of providing protection against viral challenge is present 14 days in mouse models [16,18]. A 4T1 inoculation dose ranging study was undertaken to find the maximum 4T1 cell count that would show limited tumor growth during the study and allow evaluation of the vaccine’s possible efficacy, as shown in the study design schematic in supplementary materials Figure S1. Based on the 4T1 dose ranging data shown in supplementary materials Figure S2, the challenge study was designed with one cohort receiving a 250 4T1 cell inoculation dose and the second cohort receiving 500 cells of orthotopically injected 4T1 breast cancer cells. Ten of the animals in each cohort received two doses of intraperitoneally delivered adjuvanted peptide microspheres (2.5 mg/dose), and the control mice were given two doses of blank microspheres (2.5 mg/dose, control microspheres; without peptide antigen and adjuvants) given both fourteen days before 4T1 cell inoculation, and again at the same time as orthotopic injection of 4T1cells. Subcutaneous tumor volumes were measured every three to four days with a 41-day endpoint after implantation as shown in the study design schematic in Figure 1.

**Figure 1.**
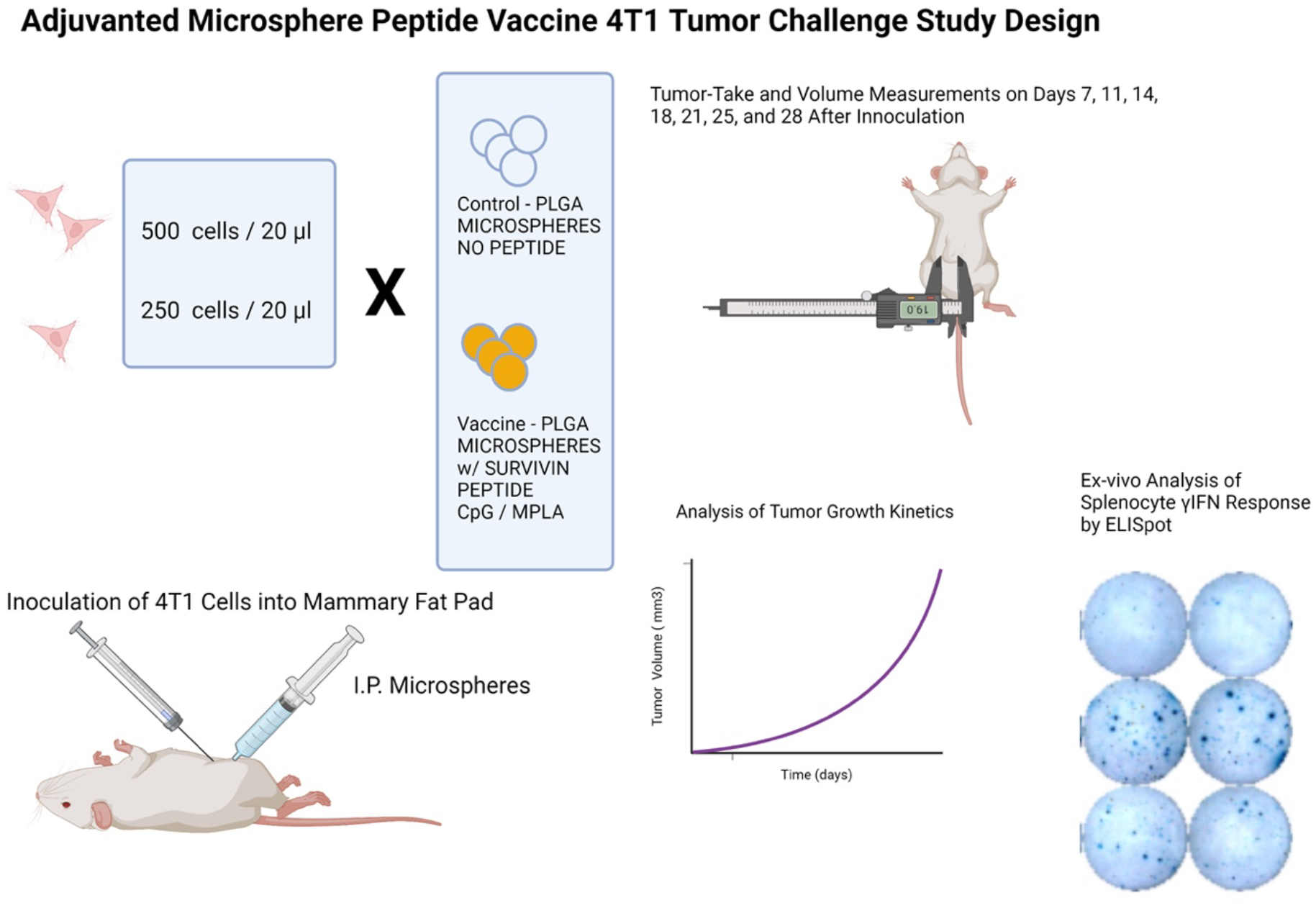
Study design schematic for a BALB/c mouse model 4T1 TNBC tumor challenge microsphere peptide vaccine efficacy study.

Mouse tumor volume was measured non-invasively every three to four days with a micrometer applied to the tumor growing in the subcutaneous space. The tumor volume (TV) was expressed in mm^3^ using the modified ellipsoid formula: TV = ½(length x width^2^)[32,33]. This data was used to calculate the tumor growth rate expressed as Δ mm^3^/day. Tumor-take frequencies were defined as the number of mice with measurable tumors on the indicated day divided by the total number of mice inoculated with 4T1 cells.

## 3. Results

Wild-type Survivin mRNA was highly expressed in all 4T1 cell line samples studied and was seen only at very low background transcription levels in the BALB/c normal mammary tissue samples (Figure 2).

**Figure 2.**
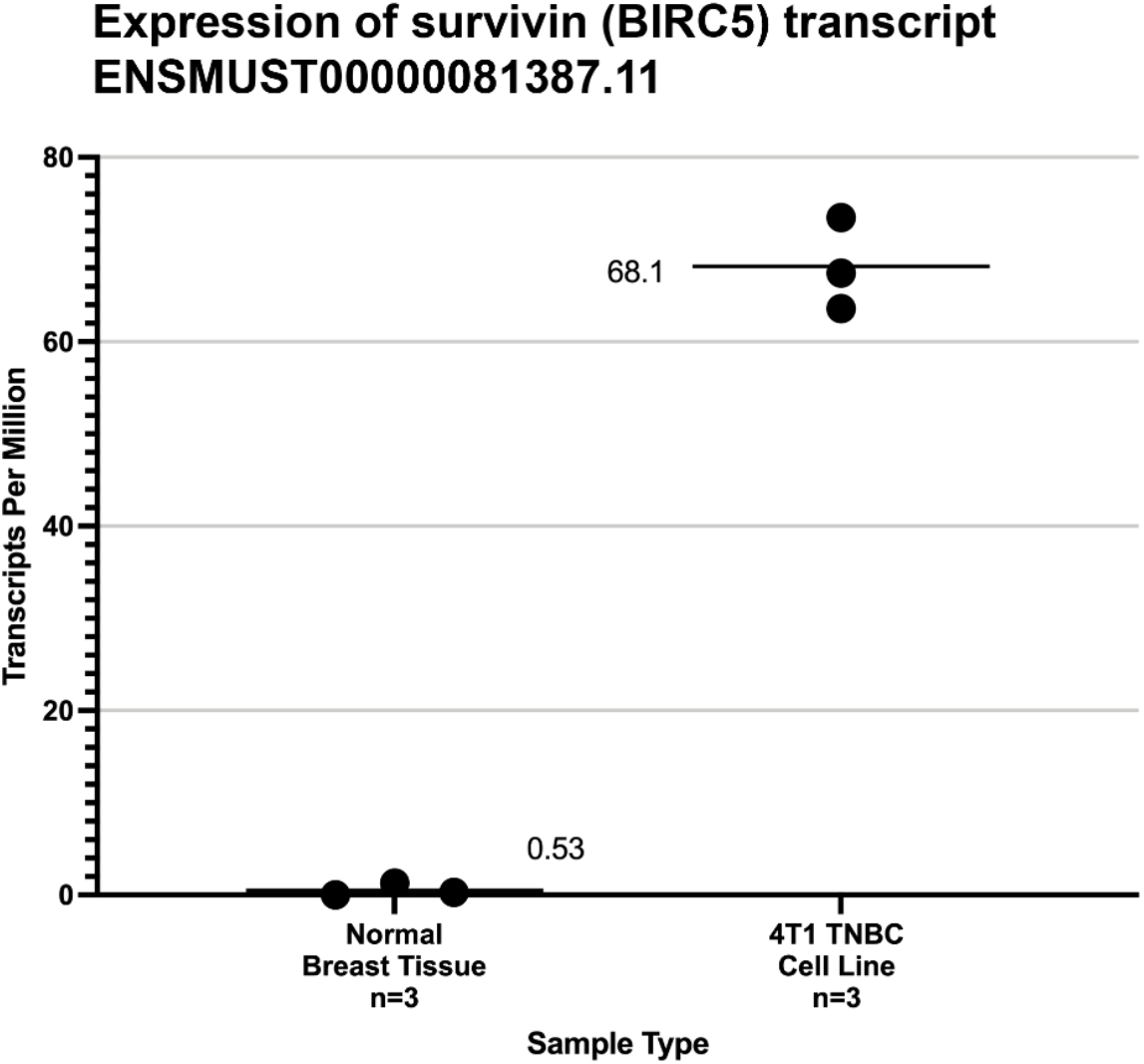
Expression level from mRNA-Seq of mouse wild-type Survivin transcript (ENST0000081387)[34] in normal breast tissue and the 4T1 cell line used in this study. The mean transcripts per millions (TPM) are shown by black bars. The significance of the difference between the average TPM found normal murine breast tissue and 4T1 cells was calculated by the method of Student.

Of the six peptides loaded into the adjuvanted microsphere formulation shown in Table 1, only QP19 (QIWQLYLKNYRIATFKNWP), produced a positive ELISpot response as shown in Figure 3. The mice who were not vaccinated did not produce a detectable response to the survivin QP19 peptide antigen. Although published studies suggested that the administered MHC class I peptide epitopes were immunogenic in BALB/c mice, we observed that only QP19 produced a T-cell response as measured by ELISpot.

**Figure 3.**
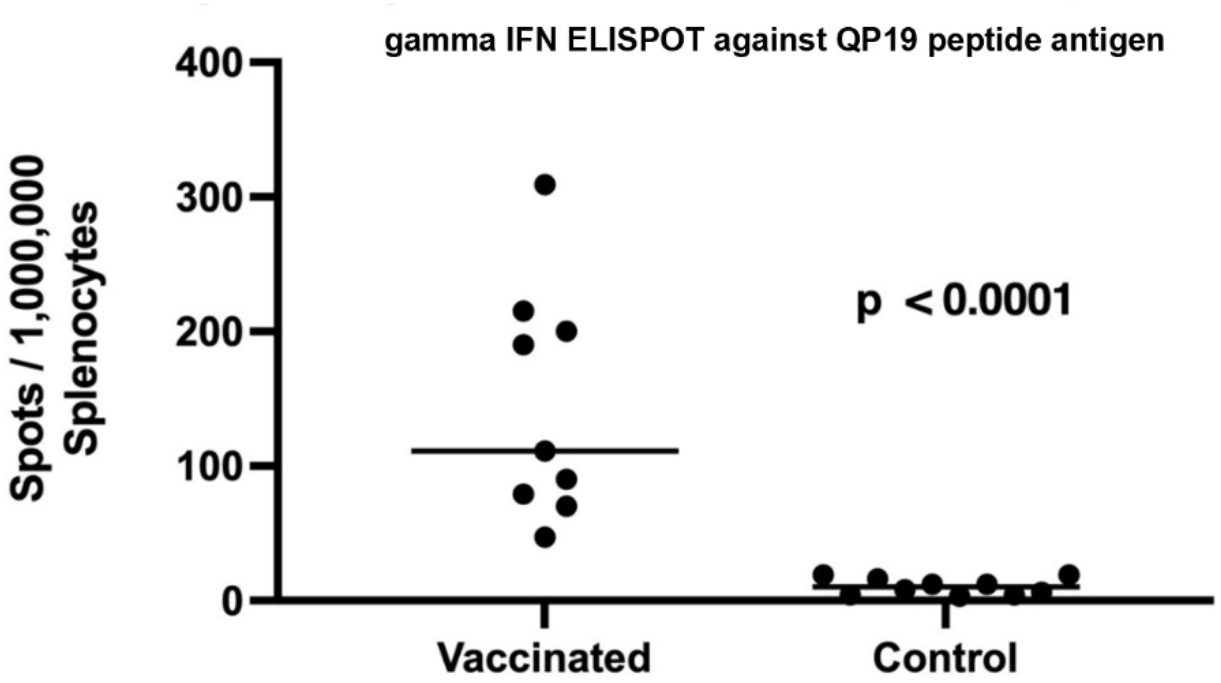
ELISpot response to QP19 in splenocytes harvested from vaccinated and unvaccinated mice that received 250 cells of 4T1. Statistical significance of the difference in the average number of ELISpots in vaccinated and control groups was determined using the unpaired, non-parametric, Mann-Whitney T-test.

The inoculation dose of 4T1 at both the 250 and 500 cell levels did not result in a tumor-take frequency of 100%, but were similar to previous reports [35] and the tumor-take frequencies measured in the dose ranging study (Figure S2). However, vaccination with survivin peptide antigens was associated with statistically significant slower primary 4T1 mammary tumor growth rates compared to tumors in control mice (Figure 4). This effect was particularly evident in the 500 4T1 cells dose group, but only at later time points / tumor volumes in the 250 4T1 cell inoculum dose group. We observed γ IFN ELISpot responses to peptide QP19 in 9/10 vaccinated mice, and noted that of these 10 mice only two mice developed growing tumors.

**Figure 4.**
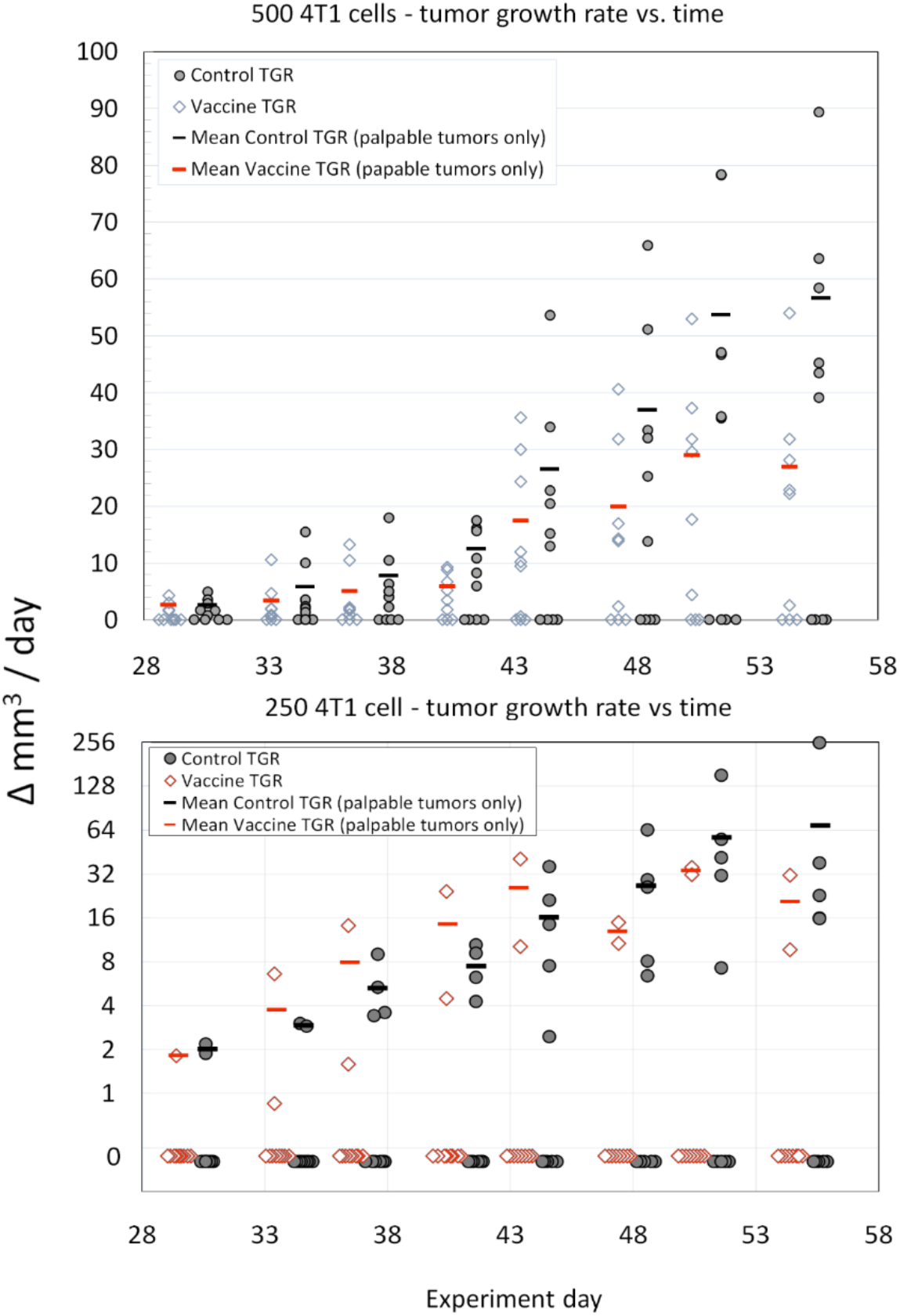
Comparison of average tumor growth rates (vaccinated versus control) for the 500 cell and 250 cell 4T1 tumor inoculation challenge groups. The sample size was n=10 for each group (Open diamonds-SVN microsphere vaccinated mice, Filled circles – control mice). The mean tumor growth rates (Red Bars – SVN microsphere vaccinated mice, Black Bars-Control mice) were calculated from growing tumors only. The statistical significance of the differences in growth rates (only non-zero data was considered) for each tumor inoculum size and treatment group (i.e., control vs. SVN peptide microsphere) was assessed using ANCOVA with Tukeys HSD and excel software according to the general linear model with time treated as a covariate. For either experiment, 500 cells or 250 cells, treatment (vaccination or control) was the significant variable (p<0.0001and p<0.009, respectively) at the 95% confidence interval.

## 4. Discussion

The targeting of survivin peptides that are HLA-matched to the host for evoking a cell-mediated immune response to kill tumor cells is of particular interest. MHC-restricted responses to peptides located on the survivin protein have been shown to elicit an immune response, including immunotherapies targeting survivin in a number of clinical trials [4]. A comprehensive list of various approaches used clinically to elicit a host immune response against survivin-expressing tumors is shown ins1s1 Table S1. A collection of HLA-restricted survivin peptide antigens identified across these various studies also raises the possibility of a broadly applicable immunotherapy for tumors that express survivin. As these studies also illustrate, eliciting a reliable, clinically significant immune response to peptide antigens is challenging.

Ensuring that the correct peptide sequence is selected and delivered effectively for T-cell expansion to occur have been obstacles to the development of safe and effective targeted immunotherapies. For example, small peptides injected on their own, even when combined with adjuvants known to enhance a T-cell response, have not been shown to trigger particularly robust T cells response [16]. As we describe here, microspheres can be manufactured that encapsulate potentially immunogenic peptides and the TLR-9 agonist CpG in a biodegradable PLGA polymer, delivered after reconstitution in a saline solution with the TLR-4 agonist MPLA to produce a cellular immune response to the administered peptide antigens as demonstrated by ELISpot [16].

In this study, we found that multiple putative peptide antigens, derived from the primary sequence of survivin and predicted to bind to MHC Class I molecules of BALB/c mice, did not elicit a detectable *ex-vivo* immune response. The QP19 peptide antigen, predicted to bind to I-A^d^/I-E^d,^ the MHC Class II molecules of BALB/c, however, was able to elicit an *ex-vivo* gamma-IFN T-cell ELISpot response and this response was associated with slower tumor growth rates. This observation suggests that protective T-cell responses were vaccine-induced and operant during tumor growth.

One possible explanation for this observation would be the presence of one or more CD8+ epitopes co-localized within the QP19 nineteen-mer, producing the observed T-cell response and anti-tumor growth activity associated with vaccination. Analysis of all possible overlapping peptides of 8-9 amino acids in length within QP19 using NetMHC and NetMHCpan found 4 potential BALB/c MHC matched peptide antigens as listed in Table S2 [26,27]. The identification of possible survivin-protective CD8+ T cell epitopes demonstrated by *ex-vivo* ELISpot response, and the formal demonstration of CD4+ T cell helper activity evoked by vaccination with QP19 await further experimentation. If such CD8+ T cell epitopes within QP19 do indeed exist, they maybe immunodominant relative to the other MHC Class I peptide SVN antigens used in these experiments. Immunodominant CD8+ T-clones may prevail over subdominant clones, masking their response. The use of anti PD-1 mAbs have been shown to promote epitope spreading in the context of anti-tumor CD8+ T cell responses [36] and certainly suggest a potential role for ICIs in SVN peptide microsphere vaccination for TNBC immunotherapy.

The use of immunotherapy for breast cancer has gained attention recently [37]. Tumor associated antigens, in contrast to neoantigens, provide the opportunity to develop immunotherapy targeting a fixed set of peptides with collective HLA restrictions predicted to provide broad population coverage that could be administered to breast cancer patients without the need for patient-specific tumor gene sequencing and manufacturing of personalized immunotherapy. Targeted T-cell immunotherapy triggering a T-cell immune response against survivin as neoadjuvant therapy has the potential to reduce recurrence after surgical excision of breast tumor tissue if the number of cells remaining after tumor debulking is low enough to result in CTL attack sufficient to blunt tumor-take and tumor growth rate. Previous studies have seen mixed efficacy with unprotected peptides used as immunotherapy[4,38]. A delivery system such as the adjuvanted microsphere encapsulation described herein, may be able to effectively deliver peptides to produce T-cell expansion against TAA targets such as survivin expressed by TNBC patients.

## Acknowledgments

A pre-print of this manuscript was posted on BioRxiv.

## Author Contributions

All authors made substantial contributions to: (1) the conception and design of the study (R.M.R., S.B., C.V.H., and P.E.H.), the acquisition of data (R.M.R., R.C. and S.B.), or the analysis and interpretation of data (P.E.H., R.M.R., S.B. and C.V.H.); and (2) drafting the article or revising it critically for important intellectual content (P.E.H., S.B., C.V.H., R.C. and R.M.R.). All authors have read and agreed to the published version of the manuscript.

## Institutional Review Board Statement

The study was conducted according to the guidelines of the Declaration of Helsinki and approved by the Institutional Animal Care and Use Committee (IACUC) of GemPharmatech Co,Ltd. (Nanjing, Jiangsu) (protocol code: SYXK (SU) 2018-0027), approval date 08/05/2019). The care and use of animals was conducted in accordance with the regulations of the Association for Assessment and Accreditation of Laboratory Animal Care (AAALAC) and the National Institutes of Health guidelines.

## Data Availability Statement

DNA and RNA-seq data can be accessed at NIH SRA under accession # PRJNA868747.

